# Reserve size and anthropogenic disturbance affect the density of an African leopard (*Panthera pardus*) meta-population

**DOI:** 10.1101/492462

**Authors:** Rasmus Worsøe Havmøller, Simone Tenan, Nikolaj Scharff, Francesco Rovero

## Abstract

Determining correlates of density for large carnivores is important to understand their ecological requirements and develop conservation strategies. Of the several earlier density studies conducted, few were done at a scale that allows inference about the correlates of density over heterogeneous landscapes. We deployed 164 camera trap stations covering ∼2500 km^2^ across five distinct habitats in the Udzungwa Mountains, Tanzania, to investigate correlates of density for a widespread and adaptable carnivore, the leopard (*Panthera pardus*). We modelled data in a capture-recapture framework, with both biotic and abiotic covariates hypothesised to influence leopard density. We found that leopard density increased with distance to protected area borders (mean±SE estimated effect = 0.44±0.20), a proxy for both protected area extent and distance from surrounding human settlements. Second, we detected a weak positive relationship between leopard density and estimated mean prey occupancy, while density was not related to habitat type. We estimated mean leopard density at 3.84 individuals/100km^2^ (95% CI = 2.53 − 5.85/100km^2^), with relatively moderate variation across habitat types. These results indicate that protected habitat extent and anthropogenic disturbance seemingly limit leopard populations more than prey abundance or habitat type. Such vulnerability is relevant to the conservation of this carnivore, which is generally considered more resilient to human disturbance than other large cats. Our findings support the notion that protected areas are important to preserve viable population of leopards, increasingly so in times of unprecedented habitat fragmentation. Protection of buffer zones smoothing the abrupt impact of human activities at reserve edges also appears of critical conservation relevance.

## Introduction

Carnivores, and large cats in particular, are not only among the most important flagship species but they also carry out critical ecosystem functions such as herbivore control, which in turn influence ecosystem health [1-3]. Yet, large cats are declining worldwide due to anthropogenic activities determining prey decline, habitat loss, unsustainable trophy hunting and direct persecution [4, 5]. Obtaining accurate density estimates for these species, and understanding the underlying factors, represents a pervasive goal in animal ecology [6]. However, this is challenging because the low abundance and elusive nature of large cats make them inherently difficult to study [2, 7].

Among the large cats, the leopard (*Panthera pardus*) has the largest distributional range in the Old World and, while it is still considered common in some areas, its range has declined by 63-75% [8]. Hunting for leopard fur and retaliatory killings for loss of livestock or human attacks, along with prey’s habitat loss, have been the major causes of such decline [8]. Leopards are highly adaptable with regards to habitat, and have been recorded in the widest range of habitat types of any Old World large feline, from mountains, rainforests and deserts to agricultural and urban areas; they are generally nocturnal, secretive in nature and have large home ranges [8-10]. Such broad adaptability in diet and habitat, along with their cryptic nature, make deciphering the relative importance of factors affecting density, such as prey abundance, habitat type, and human disturbance, particularly challenging.

Previous studies found that most often multiple correlates are associated with leopard density. In a review on carnivore abundance correlates by Carbone, Pettorelli (7), prey abundance was highlighted as the most influential factor. However, other studies found that leopard densities were not affected by prey availability but rather availability of a specific prey size category and optimal hunting habitat [11, 12]. Protected area size is another commonly assumed predictor of large carnivore densities and likelihood of their long-term persistence [13, 14]. A study conducted in South Africa addressed edge and disturbance effects on leopard abundance, and found declining density from the core of a protected area to the surrounding, unprotected landscape [15]. A study on leopards from Thailand showed a negative correlation between habitat use, with avoidance of areas with high human activity, and proximity to trafficked roads [16]. In contrast, direct anthropogenic disturbance due to encroachment into a protected area did not appear to influence a leopard population in Nepal [17], and in South Africa some leopard populations had higher densities in non-protected areas [18, 19]. In India leopards have adapted to agriculture-dominated landscapes where they occur in relatively high densities [9].

Telemetry information has been commonly used to study resource selection (e.g. [20]). However, while telemetry typically generates fine scale spatial data for a few individuals, spatial capture-recapture (SCR) sampling using camera traps generates sparse information on several-to-many individuals [21], allowing testing of explicit hypotheses on correlates of density and space use [22]. Despite the vast potential of this approach, to our knowledge there are only a dozen studies that applied robust SCR analyses to leopard density estimation; in addition, a high proportion of these studies have been performed within a single habitat type [9, 19, 23-29], while other studies have addressed differences in density between protected and non-protected areas [18, 30]. Yet the vast majority of leopard density studies have not embraced the potential of SCR analyses by incorporating drivers of species density and detectability, other than those inbuilt in the various programmes that exists.

Here, we used camera trapping across an area of ∼2500 km^2^ to estimate the density of a leopard population inhabiting a heterogeneous landscape in Tanzania, the Udzungwa Mountains. This area is a mosaic of forest blocks interspersed with drier habitats and surrounded by settled and intensively farmed areas, hence it represents a relevant landscape to study factors affecting leopard density. We used a stratified population model in a spatially-explicit capture-mark-recapture framework [31] to test hypotheses on natural and anthropogenic factors driving leopard density at the landscape scale. Specifically, we aimed to determine the effects of habitat type, abundance of potential prey, distance to water source, distance to settlements and extent of protected habitat on leopard density.

## Material and Methods

### Study area

The Udzungwa Mountains of south-central Tanzania (centred on 7°46’ S, 36°43’ E; elevation 285-2600 m asl, 16,000 km^2^) are part of the Eastern Arc Mountains, a renowned biodiversity hotspot [32, 33]. The Udzungwas consist of closed forest blocks interspersed with drier habitats [34]. It is surrounded by subsistence farming to the north, west and south, and high intensity sugar cane farming to the east, without natural habitat connectivity to adjacent protected areas [35]. Within Africa, the area is known for its outstanding levels of mammalian richness and endemism [32, 36].

The northern portion of the Udzungwas is efficiently protected by the Udzungwa Mountains National Park (UMNP; 1990 km^2^). This is surrounded to the south and west by the Kilombero Nature Reserve (1345 km^2^) administered by Tanzania Forest Service, and receives less in situ protection. A strip of agriculture, in some places just 5 km wide, separates the Udzungwa Mountains National Park (UMNP) from the Selous Game Reserve to the east (Fig 1).

**Figure 1.**
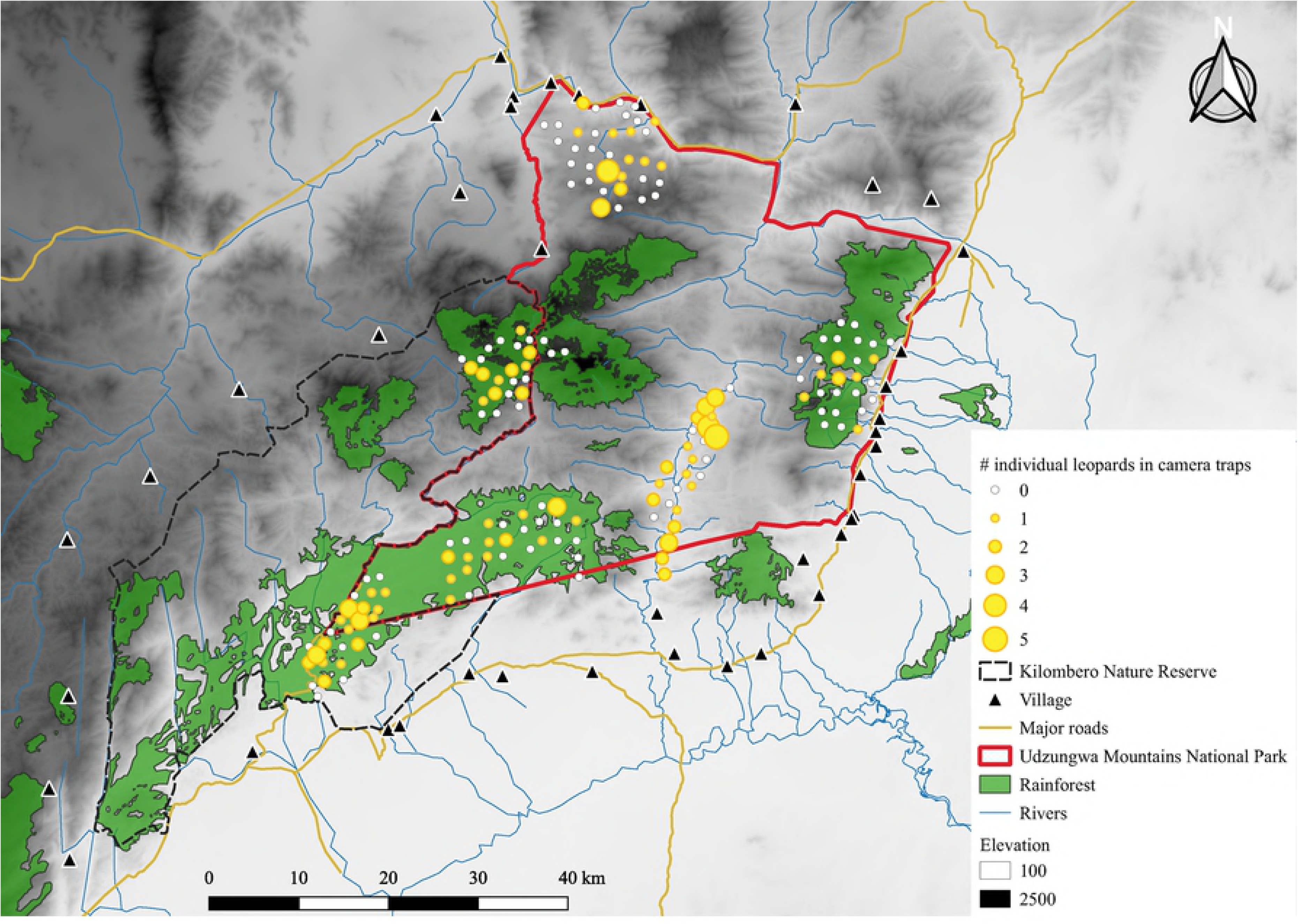
Map of the study area in the Udzungwa Mountains of south-central Tanzania. Six camera trap arrays were placed in the five major habitat types of the Udzungwa Mountains National Park and Kilombero Nature Reserve to detect leopards (*Panthera pardus*). Camera trap sites are indicated by yellow dot, with dot size indicating number of camera trap captures of leopards. Forested areas are indicated in green. Open habitat only has the elevation background. Map modified from Scharff, Rovero (37).

We placed camera trap arrays in the five, major habitat types (Fig 1; S1 Table): (1) Lowland Afrotropical rainforest in the southern UMNP (300-800 m a.s.l.). (2) Dry grassy *Acacia-Commiphora* woodlands in the northern UMNP, buffered by dry baobab woodlands at low elevation and grasslands at high elevation (500-1900 m). (3) Grassy Miombo woodlands in the central valleys of the UMNP (300-500m asl). (4) Ndundulu forest, a block of Afromontane forest west of UMNP in the Kilombero Nature Reserve (1400-2200 m). (5) Mwanihana forest, a rainforest escarpment in the eastern part of UMNP (300-2100 m).

### Camera trapping

Six camera trap arrays covering ∼2500 km^2^ were sampled sequentially to cover the five habitats. Each array consisted of 25-34 pairs of traps (Fig 1) and was sampled once each. We sampled in the dry season from August to December 2013 and from June to December 2014. Each station of paired traps operated for an average of 31 days (minimum 12, maximum 49 days; Table 1).

**Table 1.**
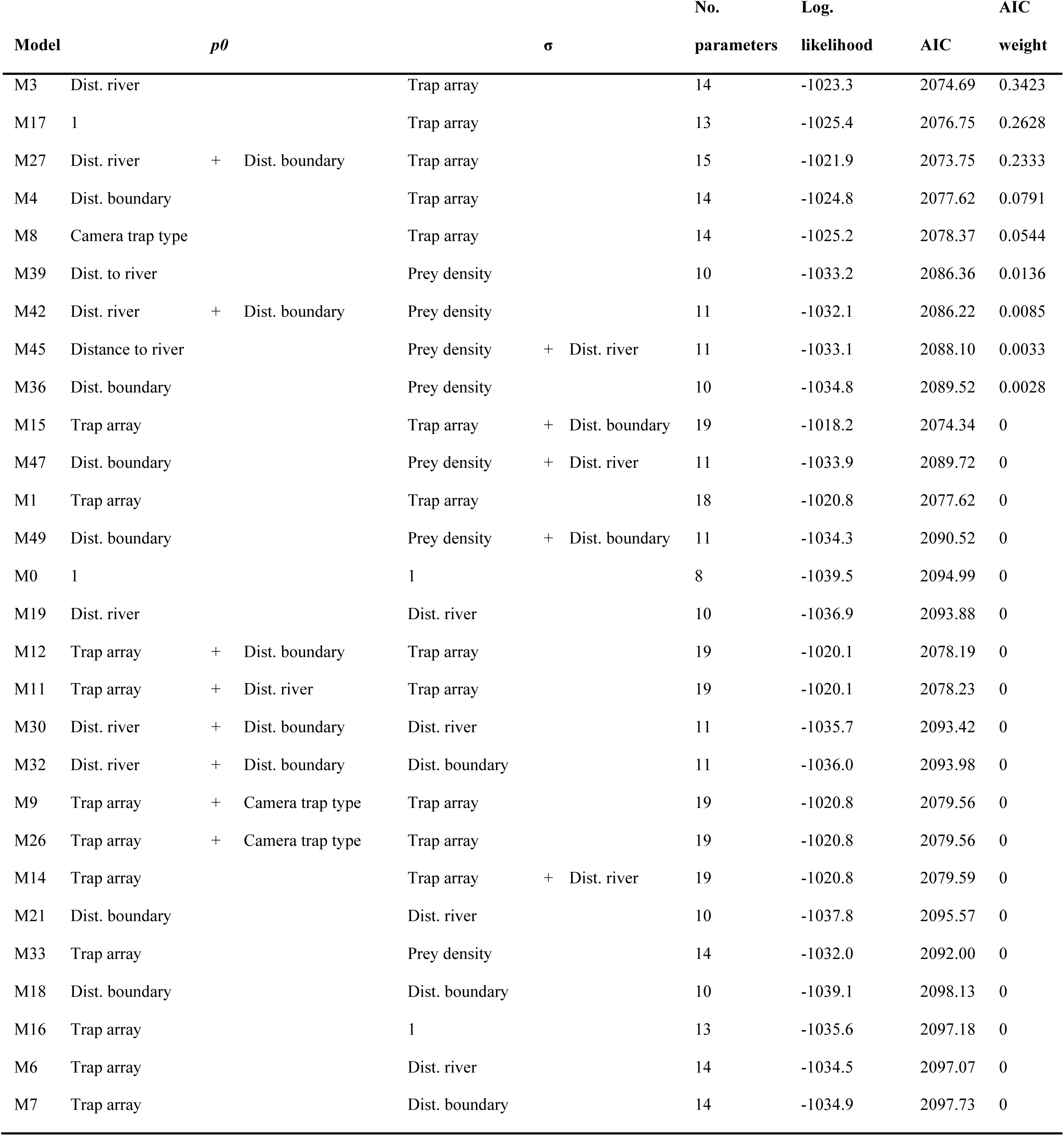
Summary of the encounter model selection for the 28 competing models.

Camera traps were set following a protocol designed for studying leopards in African forests [38]. The average trap spacing was 1.6 km, following a regular spaced grid placed randomly over the study sites. At each camera trap site, the paired cameras were placed 3-4 m from the centre of an animal trail or track, facing each other at 30-40 cm above ground level. At least one camera-trap per station had a white Xenon flash Cuddeback Ambush (Cuddeback^®^ Non Typical Inc., USA) and in 87 of 164 stations, one camera consisted of an infrared camera UOVision 565HD IR+ (UOVision Technology, Shenzhen, China) set on 15-second video recording mode.

Leopards were identified by their unique spot-patterns across their body [39] by two independent observers (RWH and FR). Only individuals deemed adult and consistently captured alone were used for subsequent analyses, to avoid non-independence of individual activity centres, e.g. juveniles [22].

### Covariates of leopard density

To model leopard density, we derived the following set of covariates across the areas covered by the six trap arrays. We first used Landsat TM and ETM+ satellite imagery to derive at 500 m resolution (1) distance from each cell centroid to the nearest river, (2) distance to the nearest protected area boundary (national park or nature reserve depending on arrays, see Fig 1). Elevation was recorded at each camera trap site using a Garmin GPS. Distance to protected area boundary correlated positively (r=0.65) with distance to the nearest human settlement from each camera trap, thus to avoid collinearity we only used distance to reserve border and considered it a proxy of both anthropogenic disturbance and extent of protected habitat. The 500 m resolution chosen for the covariates corresponds to the resolution of the state-space adopted in the spatial capture-recapture models (see below), which we defined after testing a range of resolution values that yielded stable parameter estimates and reasonable computational time. We also derived (4) an index of prey abundance as the array-specific mean estimated occupancy probability of 18 ground dwelling mammals detected by the camera traps [40]. These species were assumed to be potential leopard prey based on dietary studies [41]. In addition, 12 of these species were confirmed to be leopard prey in Udzungwa through DNA analysis of leopard scats (Havmøller (40); S2 Table). We estimated mean and array-specific occupancy probabilities (S3 Table) for the pool of potential prey by fitting a multi-species occupancy model [42] to prey species’ detection/nondetection data. This modelling approach accounts for imperfect detection and solves the ambiguity between species absence and non-detection. We therefore considered occupancy a better state variable for prey abundance than a crude index of captures, as this likely underestimate true abundance due to false negatives. In addition, as we set camera traps to target leopards, detectability of other mammals across sites may vary largely among species, likely resulting in variably biased detection rates; hence we considered it especially critical to use a state variable of abundance that is corrected by detectability [43]. We designed our community occupancy model to estimate array-specific mean occupancy values for the pool of prey species, as we assumed that the different habitat types sampled by arrays represent a relevant correlate of variation in the ‘abundance’ of prey species across the study area. However, given that we only had information on prey species at camera trap sites, we could not realistically model prey occupancy across the state-space. Specifically, we modelled the presence/absence *z*_*i,j*_ of species *i* at sites *j* as a Bernoulli trial with array-specific (*a*) occupancy probability *ψ*_*i,a(j)*_: *z*_*i,j*_ *∼ Bern*(*ψ*_*i,a(j)*_). We constrained the species-specific parameters (i.e., the heterogeneity in occupancy and detection probability) by the assumption of a common normal prior distribution for their logits. For occupancy, we considered an array-specific hyperparameter: logit(*ψ*_*i,a(j)*_) *= β*_*a*_ with *β*_*a*_ *∼ Normal*(*µ*_*ψ,a*_, *σ*_*ψ*_), where *µ*_*ψ,a*_ is the mean (community) occupancy of prey species in each array, and *σ*_*ψ*_ is the standard deviation. We organized daily detections into a species by sites matrix, with elements *y*_*i,j*_, and modelled detections as *y*_*i,j*_ *∼ Bin*(*k*_*j*_, *p*_*i,j*_\**z*_*i,j*_) where *k*_*j*_ are the sampling occasions per site and *p*_*i,j*_ is the detection probability. As we were not interested in modelling array-specific detectability, we modelled detection probability as logit(*p*_*i,j*_) = α_i_ with α_i_ *∼ Normal*(*µ*_*p*_, *σ*_*p*_), where *µ*_*p*_ is the mean (community) detectability of prey species and *σ*_*p*_ is the standard deviation. We fitted the model using a Bayesian formulation, the Markov chain Monte Carlo, implemented using the program JAGS [44] and executed from R [45]. The model code is provided in S1 Appendix. Finally, given that leopard density resulted associated significantly with the distance to reserve border (see Results), we also checked whether our prey abundance index may also be associated with this variable, hence potentially confounding the interpretation of effects on leopard density. We therefore ran a second prey occupancy model where the linear predictor for occupancy included an effect of distance to reserve border on array-specific prey occupancy. We found that for all arrays prey occupancy was not significantly associated with this covariate, hence we could exclude that the effect of reserve border on density may also be related to collinear variation in prey occupancy (see also Discussion).

### Leopard density estimation

We used spatial capture-recapture (SCR) models [22] to account for animal movement in density estimation, regarding array-specific data as samples of independent populations. This assumption is supported by the absence of individuals recorded in more than one trap array. SCR models allow study of the distribution of individuals (i.e. density) while accounting for encounter probability (p) that declines with distance between an individual activity centre (s) and a detector (x). We used a half-normal encounter model where detectability p is a function of the baseline encounter probability (p_0_) and the spatial scale parameter σ, which determines how encounter probability decreases with an increase in the distance between trap j and activity centre s_i_.

Both homogeneous and inhomogeneous point process models were fitted to study the distribution of individual activity centres within a defined state-space S, depending on the absence or presence, respectively, of spatially explicit covariates on density. We fitted a stratified population model [22] to data grouped by trap array, where array-specific population size was assumed as N_r_ ∼ Poisson(Λ_r_), where Λ_r_ is the expected number of activity centres in the state-space, or region, surrounding array r, with r = 1, …, R = 6. We investigated the effects of covariates (‘COV’, see previous session for details) on leopard density and detectability by testing different hypotheses. First, we defined the best structure of the encounter model by assessing the effect of (i) trap array, as a proxy of habitat type and seasonality (i.e. temporal variation in sampling), (ii) distance of trap j to the nearest river, (iii) distance to reserve boundary, and (iv) camera trap type on the baseline encounter probability (p_0_). The same covariates, with an additional array-specific effect of prey abundance were used as competing predictors for modelling the scale parameter σ. The general formulation of the linear predictors for two parameters of the encounter model, for individual i in trap j of array r, was as follows:

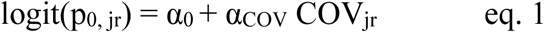

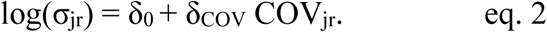

See Table 1 for a full list of competing encounter models based on plausible combination of different covariates (‘COV’). Specifically, we expected encounter rate to decline (i) with increasing distance to rivers, as waterways are frequently used as travelling routes and foraging areas [46, 47], and (ii) with increasing distance to reserve boundary, in relation to possible behavioural effects induced by an increase of anthropogenic disturbance close to the reserve boundary [12, 48]. We also expected leopards to move less (thus having smaller home range size), in dense versus open habitats, since the species has been found to prefer dense habitat for hunting and thus would have to travel smaller distances in search of optimal hunting grounds [12]. In addition, we expected leopard space usage to be (i) positively correlated with distance to the nearest river, as rivers may represent good hunting grounds [47], and (ii) positively correlated with distance from reserve boundary, where anthropogenic disturbance is higher.

We were interested in modelling density as a function of spatially-varying covariates and used a discrete representation of the state space with the centre points of each pixel g(r) (with g(r) = 1, …, G_r_) in the state-space (region) surrounding array r. The expected number of activity centres in the state-space surrounding array r was modelled in relation to (i) elevation, (ii) distance to the nearest river, (iii) distance to reserve boundary, (iv) prey abundance (occupancy probability of prey community; S3 and S4 Tables), and (vi) trap array (S1 Table). Distance to river was intended as a proxy to major traveling routes used by leopards [46, 47]. In addition, it may also indicate proximity to optimal hunting grounds. As elaborated above, we considered distance to reserve boundary a proxy for both reserve size (i.e. remoteness) and human disturbance; as settlements and farms occur right outside protected areas, we assumed human encroachment and other forms of disturbance to be more intense near boundaries [15, 16, 49]. We expected density to be negatively correlated with elevation as higher-elevation habitats are mainly in montane forests which may hold sub-optimal prey diversity and abundance and is are limited. We hypothesised a negative correlation for density with increasing distance to permanently flowing rivers as an indication of preferred hunting ground and travel routes [46]. We expected mean prey occupancy as proxy for prey abundance to be positively correlated with leopard density, matching evidence for other large carnivores and for leopards [50, 51]. We assumed leopard densities to be higher in habitats with closer, arboreal vegetation cover (montane rainforest and lowland close forest versus open woodland and wooded grassland) as these may represent more optimal foraging grounds [12]. Expected number of activity centres were modelled the in the state-space of array r in relation to the different covariates (‘COV’) as follows:

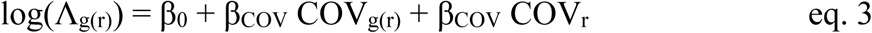

where covariates can be either spatially explicit (i.e. rasterized, ‘COV_g(r)_’) or region (i.e. survey or array) specific (‘COV_r_’). We first defined the best structure for the encounter model (27 competing models, S5 Table), while considering survey-specific densities, and then tested hypotheses on the correlates of leopard density (12 competing models, S6 Table) while keeping the best encounter structure constant. We set a 6 km buffer around each trap array based on ridged density estimates descending from 30 km and based inference on maximum likelihood estimates for leopard density using the R package ‘secr’ [31]. In order to maximize statistical power for detecting factors affecting spatial variation of leopard density on a landscape level, we used a stratified population model where data for both sexes had to be pooled. Capture histories were based on daily sampling occasions. We calculated the Akaike Information Criterion (AIC) for each candidate model and used the difference among values (ΔAIC) to rank models. We derived model averaged parameter estimates and density surfaces using model weights for those models that scored within two points of ΔAIC [52]. We also derived home range size estimates based on [22].

## Results

We accumulated a sampling effort of 5038 camera trap days and obtained 185 leopard events. Overall, 58 individuals were identified from all six surveys (median 10, minimum 5, maximum 15), excluding juveniles and sub-adults (Table 2).

**Table 2.** Capture history and effort summary from camera trapping.

| Study site | No. events | No. individuals | No. Individuals captured once | Camera traps days | Estimated leopard density / 100km<sup>2</sup> |
| --- | --- | --- | --- | --- | --- |
| Ruipa | 41 | 12 | 4 (33%) | 775 | 5.91 |
| Idete | 22 | 9 | 3 (33%) | 744 | 3.69 |
| Mbatwa | 22 | 6 | 2 (33%) | 985 | 4.46 |
| Lumemo | 53 | 15 | 4 (26%) | 887 | 4.04 |
| Ndundulu-Luhomero | 33 | 11 | 4 (36%) | 774 | 5.35 |
| Mwanihana | 14 | 5 | 1 (20%) | 873 | 4.90 |
| Total | 185 | 58 | 18 (31%) | 5038 | Average 3.84 |

Leopards were captured in 48.6% of the 164 camera trap stations. Based on AIC, the most parsimonious encounter model included an effect of distance to the nearest river on the baseline encounter probability (p_0_) and array-specific scale parameter σ (ΔAIC = 2.06; Table 1). Model-averaged estimates for the parameters of the encounter model suggest a negative relationship between the baseline encounter probability and distance of a trap to the nearest river (α_d2river_ = -0.21, -0.42 – -0.01, Table 3).

**Table 3.** Model-averaged maximum likelihood estimates for leopard populations. Estimates were computed from the two most supported models within two AIC units of each other. Model averaged estimates are provided for the parameters present in both models.

| Parameter | Description (scale) | Mean | SE | Lower 95% CI | Upper 95% CI |
| --- | --- | --- | --- | --- | --- |
| $\beta_0$ | Intercept of density (log) | -7.86 | 0.21 | -8.28 | -7.44 |
| $\beta_{d2boundary}$ | Effect of distance to reserve boundary on density (log) | 0.44 | 0.20 | 0.04 | 0.84 |
| $\alpha_0$ | Intercept of baseline encounter probability $p_0$ (logit) | -3.39 | 0.13 | -3.66 | -3.13 |
| $\alpha_{d2river}$ | Effect of distance to river on the baseline encounter probability $p_0$ (logit) | -0.21 | 0.11 | -0.42 | -0.01 |
| $\delta_0$ | Intercept of scale parameter $\sigma$ of the encounter model (log) | 7.54 | 0.15 | 7.24 | 7.84 |
| $\delta_{Lumemo}$ | Survey effect on the scale parameter $\sigma$ (log) | 0.27 | 0.19 | -0.09 | 0.64 |
| $\delta_{Mbatwa}$ | Survey effect on the scale parameter $\sigma$ (log) | 0.01 | 0.18 | -0.33 | 0.36 |
| $\delta_{Mwanihana}$ | Survey effect on the scale parameter $\sigma$ (log) | -0.50 | 0.21 | -0.90 | -0.09 |
| $\delta_{Ndundulu}$ | Survey effect on the scale parameter $\sigma$ (log) | -0.37 | 0.18 | -0.73 | -0.02 |
| $\delta_{Ruipa}$ | Survey effect on the scale parameter $\sigma$ (log) | 0.17 | 0.19 | -0.20 | 0.54 |

Baseline encounter probability (p_0_) thus decreased with increasing distance to rivers. Array-specific estimates of the spatial scale parameter of the half-normal encounter model (S7 Table) were used to derive array-specific estimates of 95% home range sizes, which varied from a minimum of 25 km^2^ in Mwanihana to a maximum of 115 km^2^ in Lumemo (mean 67 km^2^) (S8 Table).

Two models for density were most well supported based on AIC score, one including distance to reserve boundary, and a second with an additional effect of prey abundance (ΔAIC = 0.771; Table 4).

**Table 4.**
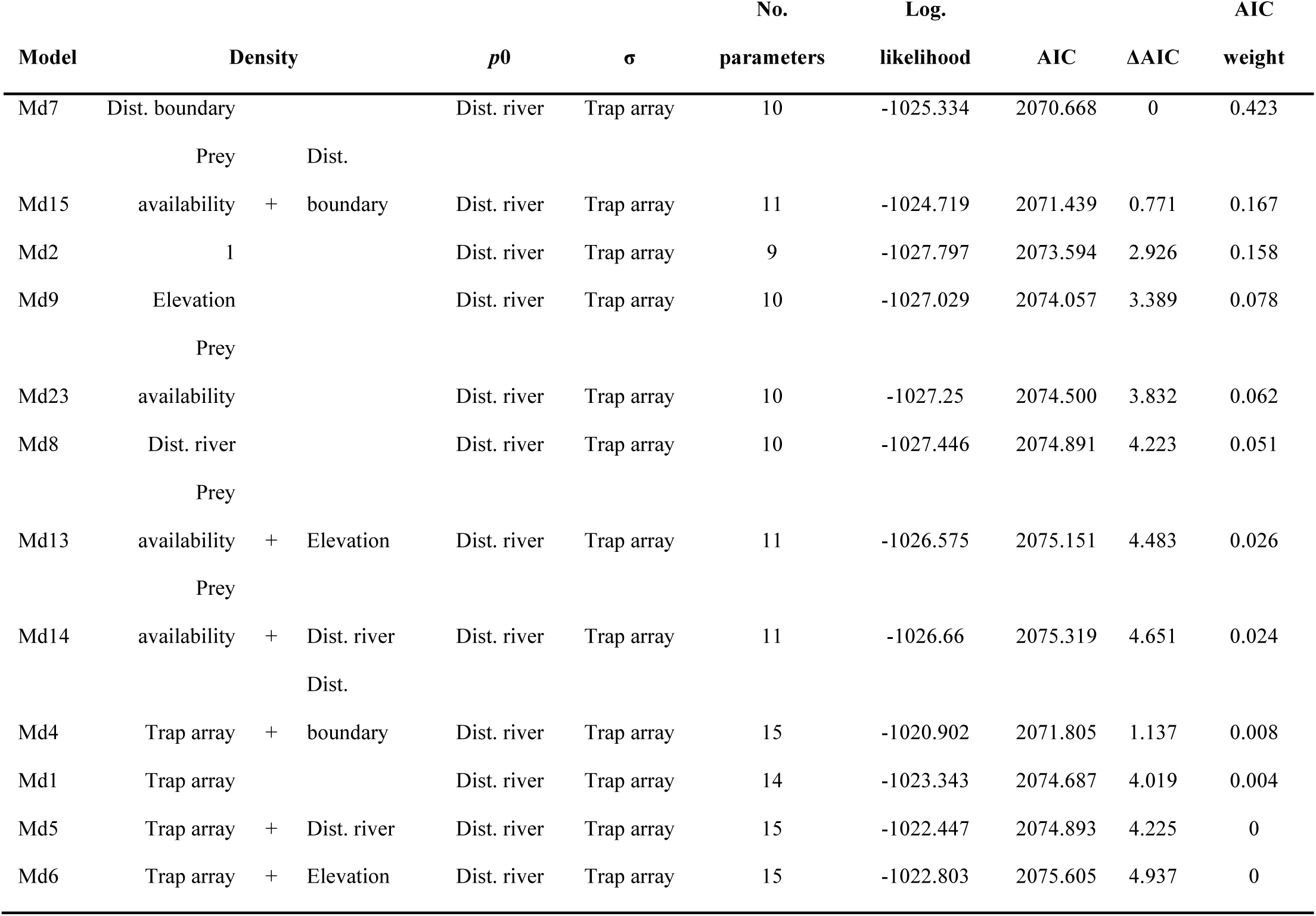
Summary for the selection of the 12 competing models of leopard density.

These two models had a cumulative AIC weight of 0.6 and suggested that leopard density was positively influenced by distance to reserve boundary, with a model-averaged β_d2boundary_ = 0.44 (95% CI = 0.04 – 0.84) (Table 3, Fig 2).

**Figure 2.**
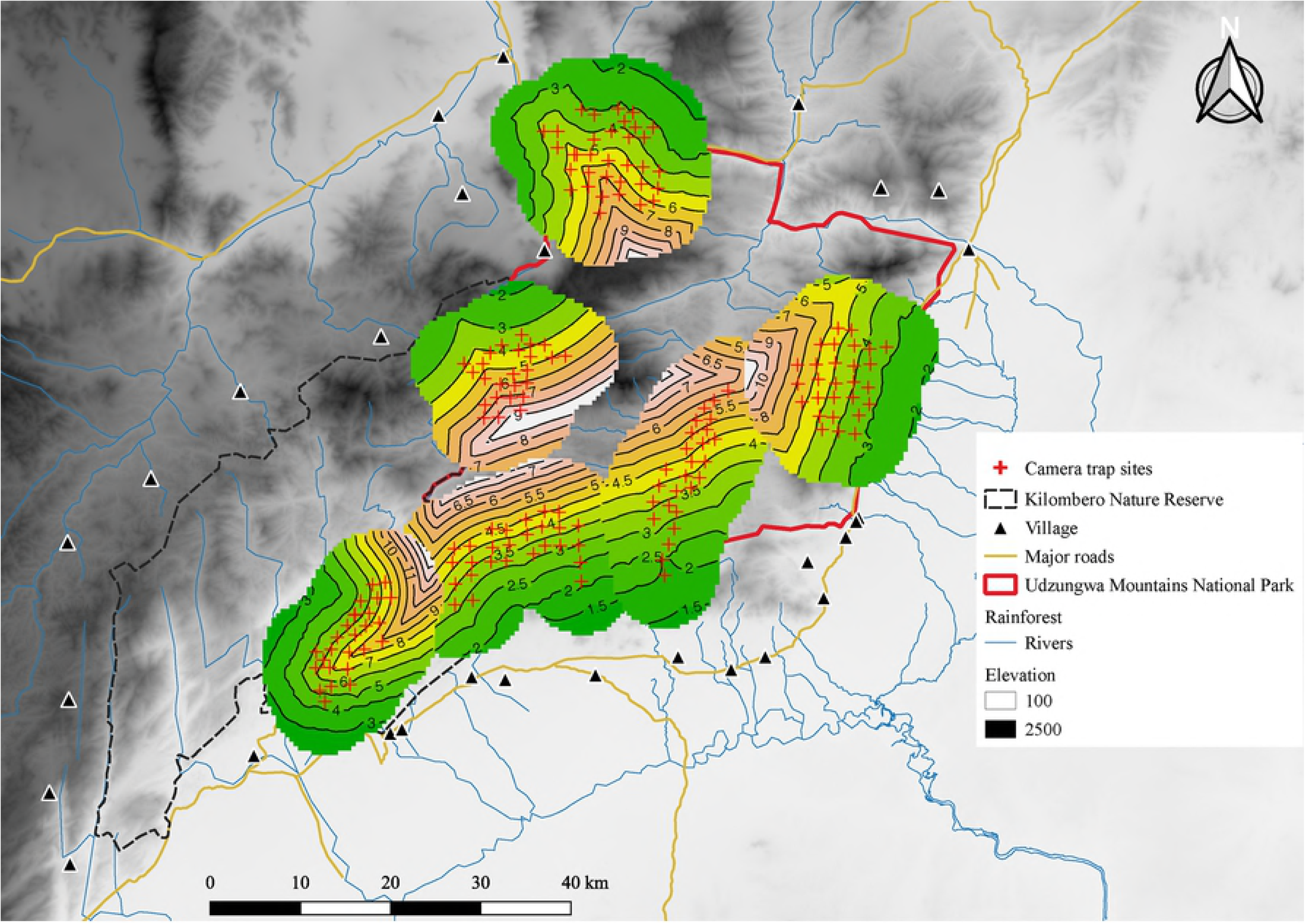
Expected leopard density (individuals/100 km^2^) in the Udzungwa mountains, Tanzania, as the predicted density surface for the state-space S superimposed over each trap array. Densities are scaled individually for each trap array with green colour indicating low densities, increasing to higher densities with warmer reddish colours. Camera trap sites are indicated as red crosses.

This effect translated into predicted densities that varied from approximately 2 individuals/100 km^2^ along the reserve border to over 8 individuals/100 km^2^ in the reserve interior (Fig 2). Average predicted density per area was lowest in Idete (3.69 leopards per 100 km^2^), largest in Ruipa (5.91/100 km^2^) and intermediate in the other areas (see Table 2). Mean density for the total area surveyed was estimated to be 3.84 individuals per 100 km^2^ (95% CI = 2.53 − 5.85/100 km^2^). As mentioned, the effect of prey abundance on density was included only in one of the two most supported models, thus not allowing the derivation of a model-averaged estimate. However, we note that despite the 95% CI of the estimated coefficient for prey occurrence included zero (mean = 0.093, 95% CI = -0.071 – 0.258; model Md15 in Table 2) the CI mainly encompassed positive values, suggesting a weak positive correlation.

## Discussion

### Correlates of leopard density at the landscape level

We analysed factors affecting the spatial variation of leopard density within a heterogeneous landscape and found that leopard density was primarily associated with distance to reserve boundary and secondarily with an index of prey abundance. We considered distance to reserve boundary a proxy for two major factors: extent of protected habitat and gradient in human disturbance. The Udzungwa Mountains National Park and adjacent Kilombero Nature Reserve form a relatively large area (3335 km^2^) of protected habitat. Concomitantly, increasing distance to reserve boundaries implies decreasing intensity of direct human disturbance from settlements [53], as indeed distance to settlements positively correlated with distance to reserve boundaries. Such heavier disturbance at reserve edges is in form of firewood collection, selective pole and timber logging, trails, poaching and charcoal production [40, 54]. Importantly, moreover, by assessing that prey abundance was not associated with distance to reserve boundary (see Methods) we could exclude that increasing leopard density away from reserve borders is mediated by an effect of increasing prey abundance.

Our findings partially mirror those from a study in South Africa, where edge effects and higher mortality rates were associated with lowered densities of leopards outside the protected area relative to inside [15]. Our result also match those of a study from Thailand, in which leopards were reported to avoid roads and areas with high human activity compared to undisturbed areas and became more diurnal when human presence became limited [48]. In a broader perspective, the magnitude of the effect of distance to reserve boundary fits the known requirement of large carnivores, for large areas of protected habitat [55]. Our results also suggest a weak positive relationship between leopard density and an index of prey abundance. We elaborate in Methods the value of this metric of occurrence that accounts for imperfect detection, a consistent issue when sampling elusive mammals in dense habitats (e.g. Dorazio et al., 2006), and therefore standardizes occupancy estimation across arrays that differ markedly in habitat type and across species. We acknowledge that a limitation of this metric is that it does not measure actual prey abundance or biomass, and it may also under-represent the full spectrum of prey species. Specifically, the species we considered did not include the arboreal primates and tree hyrax (*Dendrohyrax validus*), due to camera traps not detecting them adequately, even though for leopards occurring in the three rainforest blocks included in our landscape they seemingly represent an important prey resource [40]. Despite these limitations, we considered our metric to adequately reflect relative differences in prey abundance across our sampling arrays. Albeit we found that prey abundance had some effect on leopard density, our results in this regard partially mirror those of Balme, Hunter (12), that did not find prey abundance to be the most important factor for a population of leopards in South African savannah woodland.

We deployed six camera trap arrays covering five habitat types that differ markedly in vegetation cover, from montane to lowland rainforest, dry forest and wooded grassland (Fig 1; Table 1). However, we found that the mean den<u>s</u>ity of leopards varied little across these habitats (3.69 – 5.91 individuals per 100 km^2^), and model selection indicates these estimates do not substantially differ. Therefore, we conclude that disturbance avoidance and, secondarily, prey abundance are more important factors than habitat type for leopards in our study landscape. This conclusion fits the notion of leopards being habitat generalists [56], and in our case study their apparent flexibility in respect to habitat may also be contributed by the fact that leopards are the most abundant large carnivore in Udzungwa (spotted hyenas [*Crocuta crocuta*] occur in lower density and lions [*Panthera leo*] are only occasionally recorded [40]), thus leopards are not constrained by interactions with other large carnivores.

We found that baseline encounter probability (p0) was positively correlated with proximity to rivers, while space-usage (σ) changed with habitat type. Higher encounter probability close to waterways may be related to habitat structure, with large and frequently used trails cutting across dense vegetation that may result in optimal detection of animals by camera traps, as opposed to less dense habitats. Travelling along rivers is also known to be more energy efficient and favoured places for scent marking and hunting [38, 46, 47]. Higher abundance close to rivers may increase encounter probability if the two variables are positively correlated. However, we did not find support for a significant relationship between density and distance to river.

### Conservation implications

We considered a suite of natural and anthropogenic factors hypothesised to affect leopard densities in a complex landscape with different habitat types. We found that distance to protected area boundary, which was in turn correlated to distance to human settlements, was the single, most influential factor affecting leopard density. We also found that the importance of this factor overwhelmed the influence of prey abundance, which was retained in the second-best model, while habitat type did not have an effect. These results support the notion of high flexibility of the leopard with regards to prey and habitat preferences [41]. However, intriguingly we had no recaptures of individual leopards between the major habitat types despite their relative proximity. Our mean population density estimate of 3.84 leopards/100km^2^ appears in the mid-range when compared to densities from comparable areas in Africa, where high density estimates of 12.03/100km^2^ are known from Kenya [29] and low estimates of 0.59/100km^2^ are known from Namibia [25]. In Udzungwa, leopards are reported as extremely rare or locally extinct in the least protected parts of the Kilombero Nature Reserve [57] and in smaller and poorly protected forests in the range, such as Uzungwa Scarp [58]. Indeed recent research shows that the reserves adjacent to UMNP have much lower mammalian abundance and species richness and that this is associated with their level of protection [54].

Leopards disappearance has been attributed to direct hunting and prey depredation [59], mirroring findings from the Congo [60] and South Africa [15]. While our study shows populations in the protected area are in decent densities, the regional metapopulation could be at risk if they lose connectivity with the major adjacent ecosystems of Selous and Ruaha. Our findings carry important conservation implications, which are related to the need for maintaining large areas of continuous, well-protected habitat to preserve viable population of large carnivores [13]. This becomes even more imperative given the ever increasing habitat fragmentation that terrestrial mammals face globally [61]. The establishment of buffer zones, for example in form of Wildlife Management Areas (WMAs), i.e., areas co-managed with local communities [62], may be a feasible option for the Udzungwa landscape. WMAs would ensure greater protection along the currently abrupt edges between reserves and human settlements [63].

## Acknowledgements

We thank Roland Kays for constructive comments on an earlier version of the manuscript. We also thank the Tanzania Wildlife Research Institute and Tanzania National Parks for their collaboration and assistance and the staff of the Udzungwa Ecological Monitoring Centre for logistic support during field work. In particular, Richard Laizzer and Aloyce Mwakisoma for invaluable field assistance. NS and RWH acknowledge the Danish National Research Foundation for funding for the Center for Macroecology, Evolution and Climate (grant no. DNRF96) with co-funding from MUSE – Science Museum (Trento, Italy). RWH is supported by the Carlsberg Foundation (CF16-0310 & CF17-0539) and would also like to thank Tom Gilbert from the Section of Evolutionary Genomics for support, funding and supervision. FR and RWH would like to thank Fototrappolaggio s.r.l. for sponsoring camera traps. This research project was conducted under COSTECH permits 2013-274-NA-2013-111 and 2014-137-ER-2013-111.

